# Gait signature changes with walking speed are similar among able-bodied young adults despite persistent individual-specific differences

**DOI:** 10.1101/2024.05.01.591976

**Authors:** Taniel S. Winner, Michael C. Rosenberg, Gordon J. Berman, Trisha M. Kesar, Lena H. Ting

## Abstract

Understanding individuals’ distinct movement patterns is crucial for health, rehabilitation, and sports. Recently, we developed a machine learning-based framework to show that “gait signatures” describing the neuromechanical dynamics governing able-bodied and post-stroke gait kinematics remain individual-specific across speeds. However, we only evaluated gait signatures within a limited speed range and number of participants, using only sagittal plane (i.e., 2D) joint angles. Here we characterized changes in gait signatures across a wide range of speeds, from very slow (0.3 m/s) to exceptionally fast (above the walk-to-run transition speed) in 17 able-bodied young adults. We further assessed whether 3D kinematic and/or kinetic (ground reaction forces, joint moments, and powers) data would improve the discrimination of gait signatures. Our study showed that gait signatures remained individual-specific across walking speeds: Notably, 3D kinematic signatures achieved exceptional accuracy (99.8%, confidence interval (CI): 99.1-100%) in classifying individuals, surpassing both 2D kinematics and 3D kinetics. Moreover, participants exhibited consistent, predictable linear changes in their gait signatures across the entire speed range. These changes were associated with participants’ preferred walking speeds, balance ability, cadence, and step length. These findings support gait signatures as a tool to characterize individual differences in gait and predict speed-induced changes in gait dynamics.

## 1. Introduction

Seemingly stereotypical human behaviors such as walking and running exhibit distinct individual characteristics^[1–5]^ that stem from complex interactions of neural control, muscle activation patterns, biomechanics, sensory feedback, and the environment. Precisely because of these interconnected processes, the mechanisms underlying individuality in gait coordination across speeds remain elusive. Previously, we developed a proof-of-concept framework leveraging a recurrent neural network (RNN) model to capture individual differences in human gait dynamics based on measured kinematics^[6]^. The high-dimensional internal parameters of the trained model encode how individuals’ inter-and intra-limb gait variables progress over time. From the model’s internal activations, we constructed low-dimensional representations of individuals’ multi-joint coordination, phase-averaged across multiple strides, termed *gait signatures*. Gait signatures have broadened our understanding of human inter-joint coordination, going beyond traditional analyses focused on discrete spatiotemporal^[7]^, kinematics and kinetics^[8]^, and other derived metrics of gait coordination^[9–13]^. Thus, our prior work introduced a method that showed promise using 2D joint angles to identify individual differences in gait signatures that remain individual-specific across a restricted range of walking speeds in both able-bodied and impaired gait^[6]^. Here, we tested the ability of gait signatures, derived from different data types (3D kinematics and 3D kinetics), to discriminate individuals amidst a wide range of speeds. Additionally, we examined gait signatures in able-bodied young adults across a wide range of walking speeds to understand their speed-dependent variations. Finally, to uncover potential factors influencing speed-dependent modifications in gait signatures, we investigated whether these changes correlate with specific spatiotemporal variables and dynamic balance.

We hypothesize that individual differences in gait dynamics persist across a wide range of walking speeds. This persistence allows us to identify individuals by their gait signatures regardless of gait speed, despite the biomechanical changes required to walk at different speeds. To modulate gait speed from slow to a faster pace, individuals can employ various strategies, including taking longer steps, increasing step frequency, and decreasing the swing or stance phase durations, among others^[14–17]^. Increasing gait speed has also been associated with complex changes in joint excursions and step length asymmetry in people post-stroke^[18,19]^. However, previous research demonstrated that speed had little effect on joint-level coordination in injury-free adults^[20,21]^. Building on our previous findings^[6]^, where we demonstrated discernible individual gait signatures across a limited range of speeds, we anticipate that although gait signatures would change across a wider speed range, their individual-specific nature would be preserved.

Understanding what data are needed to differentiate gait coordination patterns between individuals may inform experimental design, equipment considerations, and future clinical translation. Our prior work^[6]^ constructed gait signatures using only sagittal-plane joint kinematics (i.e. joint angles). However, including additional kinematic and kinetic (i.e. ground reaction forces, joint moments, and powers) data may offer further insights into the individual-specificity of gait signatures. While majority of gait variability occurs in the sagittal plane, variability may also manifest in the frontal and coronal planes among certain able-bodied individuals, evident in movements like hip abduction/adduction and internal rotation^[22]^. This variability becomes more pronounced in impaired individuals, such as stroke survivors, who often adopt compensatory strategies in the frontal plane, like circumduction or pelvic hiking^[23–25]^. Furthermore, kinetic information may be important to include as kinetics might contain meaningful information not encoded in the kinematics^[25]^. For example, two people with identical kinematics and different body compositions will have different kinetics^[26]^. Alternatively, estimated joint kinematics and kinetics are mechanically related, such that kinetics may not be necessary to distinguish individuals if soft-tissue artifacts and measurement errors^[27]^ are negligible, compared to individual differences in kinematics^[28–30]^.

Given similar biomechanical constraints and unimpaired neural control in able-bodied young adults, we hypothesize that there exist common changes in gait dynamics across speeds. Numerous studies have reported many discrete gait parameters that increase with gait speed, including leg stiffness^[31]^, center-of-mass work and power^[32]^, muscle activity amplitude^[33]^, joint angle excursions^[34]^, kinetic variables^[20,35]^, and spatiotemporal variables^[14,16,17,36,37]^. Moreover, previous research has identified many relationships between specific gait characteristics and walking speed^[32]^. For example, spatiotemporal parameters demonstrate squared and cubic relationships, while kinematic outcomes exhibit a range of linear, squared and cubic relationships with normalized gait speed^[38]^. Additionally, certain kinetic parameters exhibit a linear relationship with gait speed, while others display quadratic relationships^[39]^. While common measures of gait biomechanics exhibit differential relationships with speed, we tested whether gait signatures would also exhibit a consistent relationship with speed. As gait signatures leverage continuous, synchronously measured gait data to identify a low-dimensional representation of gait dynamics, they may provide a more comprehensive representation of how gait coordination changes with speed. This approach may shed light on the underlying organization of gait variables, enriching our understanding of how gait dynamics change across speeds.

Understanding whether changes in gait signatures across different speeds correlate with changes in discrete spatiotemporal variables and dynamic balance ability would highlight potential factors impacting how people modulate coordination across speeds. Studies show that walking at extremely slow speeds disrupts individuals’ natural momentum and coordination^[40]^ and biomechanical strategies^[41]^. We hypothesize that an individual’s capacity to modulate their gait with speed is contingent upon underlying factors inherent to their sensorimotor system and functional capacity. We predict that individuals with better balance ability may be able to adapt more flexibly (exhibit less change in their signature) to walking at extremely slow or fast treadmill speeds than those with lower balance ability.

This study assesses whether able-bodied young adults’ gait signature remain individual-specific walking speeds ranging from extremely slow (0.3m/s) to exceptionally fast (above the empirical walk-to-run transition). First, we determined the optimal number of speed trials per individual required to a train a linear support vector machine classifier effectively, enabling accurate individual identification across various speeds for each data type. Next, we characterized how the data type used to train the gait signatures model (2D kinematics, 3D kinematics, 3D kinetics and their combination) impacted the ability to identify individuals based on their gait signatures using a support vector machine classification task. Thirdly, we characterized the extent to which individuals exhibited consistent changes in gait signatures with speed. Lastly, we determined whether changes in gait signatures with speed were associated with changes biomechanical variables and individuals’ dynamic balance ability.

## 2. Materials and methods

### 2.1 Ethics statement

All participants provided written informed consent prior to study participation, and the study protocol was approved by the Emory University Institutional Review Board.

### 2.2 Human subject participants

Seventeen young able-bodied adult individuals participated in this study (8 men, 9 women; mean ± s.d. age = 27.9 ± 4.5 years, height = 1.7 ± 0.1 m, body mass = 77.0 ± 21.8 kg). Participants reported no history of injury or pain in the last 3 months.

### 2.3 Experimental design

To test the effect of a wide range of speeds, participants completed 60-second trials at 9 different speeds ranging from extremely slow to extremely fast speeds. The speed conditions were implemented in 2 phases, and within each phase speeds were assessed in random order. Phase 1 was always implemented first and consisted of 6 speed conditions ranging from the fixed extreme slow speed of 0.3 m/s to each participant’s fastest comfortable safe speed determined on the treadmill. Phase 2 was implemented second and consisted of the 3 remaining speed conditions (92.5%, 100.0%, and 107.5% of the empirical walk-to-run transition speed). Participants were advised to take a seated or standing rest break for 1-2 minutes as needed, and if they experienced fatigue or pain following a gait trial. The 9 speeds evaluated for each participant are outlined below as follows:

#### Phase 1 speed conditions

1. <u>Fixed extreme slow</u>: a very slow fixed speed of 0.3 m/s was selected to match the speed of low functioning stroke survivors.
2. <u>Slow overground-derived</u>: participants were instructed to walk at a very slow pace overground (instruction: “walk as if leisurely strolling in a beautiful park”) along a flat, indoor marked 29.9-foot (9.11-meter) path in a controlled lab setting. Three trials were performed and the average speed for this condition was calculated for each participant.
3. <u>Self-selected treadmill-derived</u>: The treadmill was initiated at ∼1 m/s and participants were instructed to indicate whether to increase or decrease the speed until they reached a speed that was representative of their natural or comfortable walking speed.
4. <u>Self-selected overground-derived</u>: participants were instructed to walk at their natural self-selected pace overground (instruction: “walk at a pace that is natural for you to travel from point A to B”) along a flat, smooth, marked 29.9-foot path in a controlled lab setting. Speed was determined as the average speed from three trials.
5. <u>Intermediate calculated:</u> The speed halfway in between each participant’s self-selected overground-derived and the fast treadmill-derived speed was calculated.
6. <u>Fast treadmill-derived</u>: The treadmill was initiated at ∼1 m/s and participants were instructed to indicate whether to increase or decrease the speed until they reached a speed that was representative of a fast-walking speed (instruction: “walk as if you are running late for a very important event”).

#### Phase 2 speed conditions

7. <u>92.5% of the walk-to-run transition speed</u>: 7.5% below the determined walk-to-run transition speed was calculated.
8. <u>Walk-to-run transition speed</u>: The walk-to-run transition speed was determined before randomizing the 3 speed conditions in phase 2. The treadmill was initiated around each individuals’ determined fast walking speed, and participants were instructed to indicate whether to increase or decrease the speed until they reached a speed that felt they could no longer walk and needed to start running. Treadmill speeds were varied by 0.05m/s - 0.13m/s at a time. Once participants identified a preferred transition speed, the treadmill speed was increased beyond, then decreased below, that speed to encourage exploration of each speed. Participants were instructed to try walking and running at each speed (∼20 seconds), if possible. Next, participants were again asked to identify the speed beyond which they prefer to run and below which they prefer to walk. This process was repeated until participants selected two consecutive speeds within 0.05 m/s of each other. The recorded speed was the one they settled on during the process. This approach is similar to another study^[42]^ however, we did not mandate rest periods.
9. <u>107.5% of the walk-to-run transition speed</u>: 7.5% above the determined walk-to-run transition speed was calculated.

### 2.4 Data acquisition and processing

We used 3D motion capture to obtain continuous walking data from participants. Reflective markers were attached to participants’ trunk, pelvis, and bilateral shank, thigh, and foot segments^[43]^. We collected marker position data while participants walked on a split-belt instrumented treadmill (Bertec Corp., Ohio, USA) using a 7-camera motion analysis system (Vicon Motion Systems, Ltd., UK). Marker data were collected at 100Hz, and synchronous ground reaction forces were recorded at 2000 Hz and were down sampled to 100Hz using previously established techniques^[43–46]^. Raw marker position data were labeled and gap-filled. Marker trajectories and ground reaction force raw analog data were low-pass filtered at 6 and 30 Hz in Visual 3D (C-Motion Inc., Maryland, USA). Gait events (bilateral heel contact and toe-off) were determined using a 20N vertical GRF cutoff; 3D kinematics and kinetics were calculated in Visual 3D.

To describe the 3D kinematics of each individual, we measured 18 continuous variables from the motion capture data; bilateral hip, knee, and ankle joint angles each in the sagittal, frontal, and transverse planes. Two-dimensional kinematics consisted of 6 continuous features, comprising bilateral hip, knee, and ankle joint angles in the sagittal plane only. 3D kinetics consisted of 42 total features-bilateral ground reaction forces normalized to body weight, ankle, knee and hip moments and powers each in the sagittal, frontal, and transverse planes. A combination of all the data consisted of 60 variables. sagittal-plane hip, knee, and ankle joint kinematics. Three speed trials were omitted (2 for participant YA04 and 1 for participant YA06) due to technical errors during data collection, resulting in a reduction of the full trial data set from N = 153 to N = 150. The excluded trials corresponded to YA04 and YA06’s 7.5% below walk-to-run transition speed, and YA4’s 7.5% above walk-to-run transition speed. To visualize representative traces of 3D kinematics and kinetics across different speeds, refer to (Fig. 1). One minute of continuous 3D motion capture treadmill walking data were collected from 17 able-bodied young adults across 9 different speed conditions (Fig. 2a). Continuous gait data from all individuals and speed conditions were extracted (Fig. 2b).

**Figure 1:**
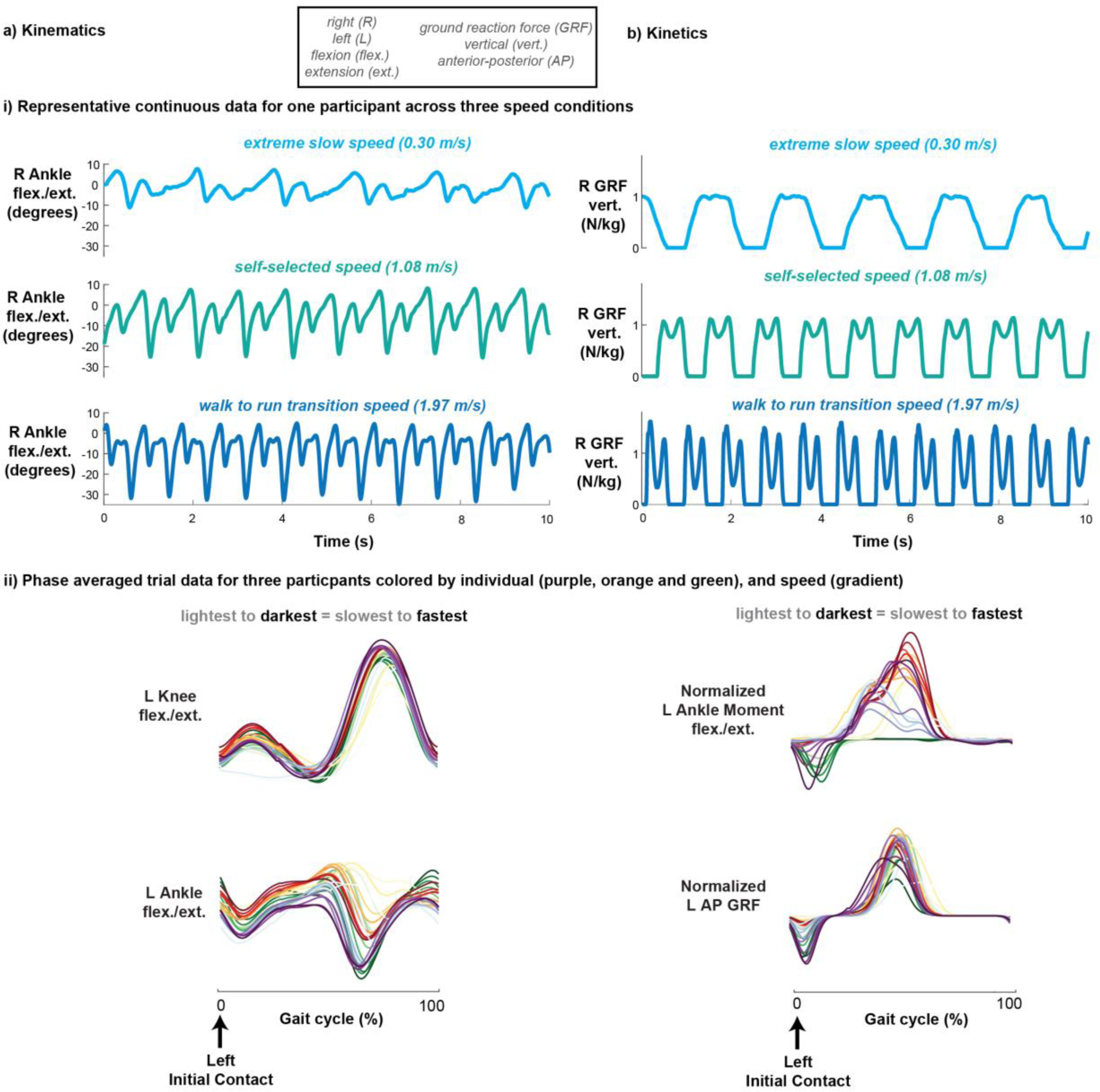
Visualization of representative a) kinematic and b) kinetic treadmill walking data. Representative continuous right angle flexion/extension (a-i) and right vertical ground reaction force (b-i) for one participant across three speed conditions: extreme slow, self-selected, and extreme fast (walk to run transition). Representative phase-averaged left knee flexion/extension (a-ii, top), left ankle flexion/extension (a-ii, bottom), normalized left ankle moment flexion/extension (b-ii, top), and normalized left anterior-posterior ground reaction force (b-ii, bottom) data for three participants colored by individual (color) and speed (gradient).

**Figure 2:**
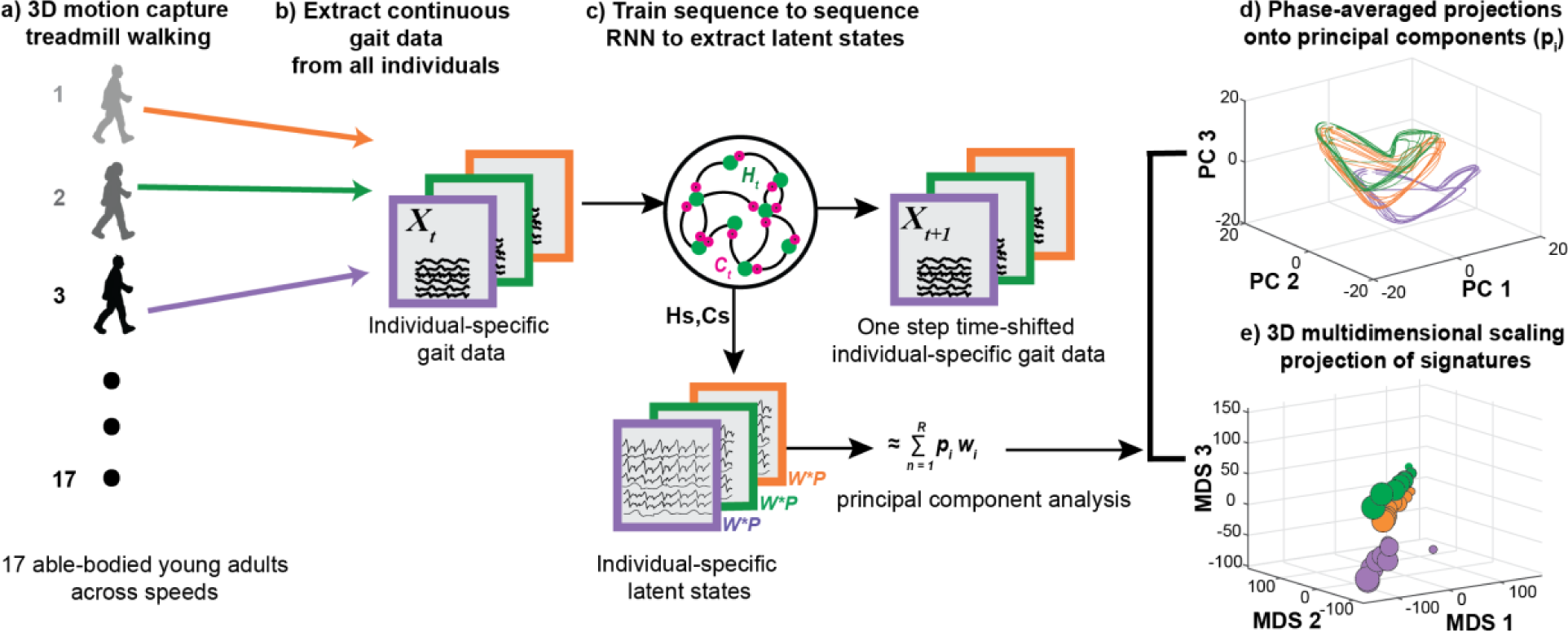
Pipeline of the gait signatures framework and outcomes. a) 3D motion capture data from 17 able-bodied young adults walking on a treadmill across 9 speeds each was conducted. b) Continuous timeseries kinematics and kinetics were extracted from all trials. c) A sequence-to-sequence RNN was trained using subsets of the data recorded in (b), and individual-specific gait signatures were extracted for all individuals’ trials. d) Principal component analysis was applied to reduce the dimensionality of the high dimensional latent states, each trial was phase-averaged, and the first 3 dominant PCs visualized as 3D loops. e) 3D projections of low-dimensional gait signatures using multidimensional scaling reveal individual-specific gait signatures among 3 representative able-bodied young adults.

### 2.5 Generating gait signatures

To create gait signatures, we utilized the RNN framework described previously^[6]^, where continuous, multi-joint kinematics from multiple individuals and speeds were used to train an RNN model. The RNN model architecture consisted of a single input layer, a hidden layer with 512 long short-term memory (LSTM) activation units, and an output layer. The model was trained using an Adam optimizer^[47]^ in a sequence-to-sequence manner to predict one-step time shifted output kinematics (Fig. 2c). To prevent overfitting on this larger dataset of stereotypical able-bodied gait patterns, we made the following modifications to our prior framework^[6]^. We lowered the learning rate of the Adam optimizer from 0.001 in to 0.0001, added a drop out layer after the hidden layer with 20% drop out rate, and added a kernel L2 regularization (regularization strength of 0.01). Additionally, the trials in this able-bodied young adult dataset were 60 seconds long vs. the 15 second trials used previously. During training, our data were batched according to the number of total trials (N = 150). and the RNN was trained on all individuals trials to extract individual-specific latent states of the RNN which represent individuals’ gait dynamics (Fig. 2c, Individual-specific latent states).

The RNN latent states (Fig. 2c, Individual-specific latent states), were extracted for all individuals’ trials, reduced in dimension using and principal component analysis and phase-averaged^[48]^ to generate gait signatures. Gait signatures can be visualized as looped representations of specified principal component (PC) projections (Fig. 2d) and 3D multi-dimensional scaling projections (Fig. 2e). We trained four individual RNN models, each with a different input data type (2D kinematics, 3D kinematics, 3D kinetics and a combination of the data). Gait signatures were generated for each model RNN model separately.

### 2.6 Determining the variance explained in the original features by the principal components of the gait signatures model

To attain the variance explained in the original features by each principal component (PC) of the RNN latent spaces, we used each trained RNN model (Fig. 3a, 1. Train RNN), extracted the model internal activations (weights), performed PCA on them, and identified the weights corresponding to each PC (Fig. 3a, 2. Isolate weights corresponding to each PC). The model weights were updated to a new model based on the top N principal components and used to generate reconstructed data for the provided internal states (Fig. 3a, 3. Reconstruct data for isolated PCs). The coefficient of determination (R^2^) was calculated between the reconstructed data and the measured data (Fig. 3a, 4. Evaluate the R^2^ between measured and reconstructed data for each PC). Eigenvalue plots of the cumulative variance explained by each PC of the gait signature (expressed as a percentage) were constructed for each data type (Fig. 3b).

**Figure 3:**
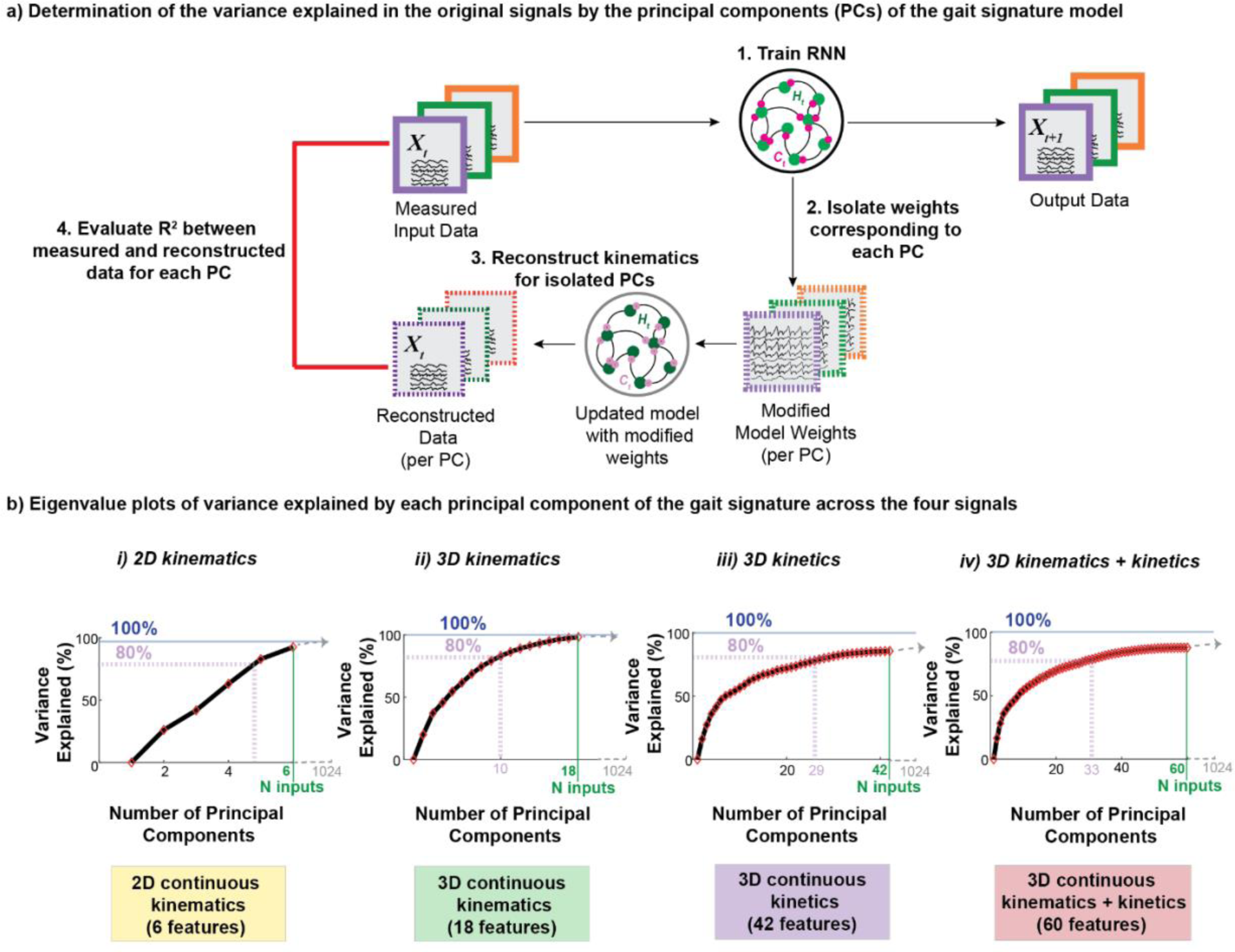
Variance explained in the original signals by each of the principal components of the extracted dynamics. a) To determine the variance explained in the original data by the dynamics: 1) the RNN model was trained, 2) the trained model weights (internal activations) were extracted and the dynamics corresponding to each PC was isolated. 3) For each PC, the original signals were reconstructed across participants. 4) The coefficient of determination was calculated between the measured input signals and the reconstructed signals for each PC. These values were used to construct eigenvalue plots for each signal type. b) Eigenvalue plots of the cumulative variance explained by increasing number of principal components of the gait signature for each of the four signal types.

To determine the number of principal components to retain, the elbow of the eigenvalue plot is usually considered sufficient^[49]^. However, since our eigenvalue plots represent the variance explained by PCs of the original data (not of the high-dimensional gait signatures i.e., internal activations), we determined a reasonable threshold of 80% to account for most of the variance explained in the original model input data.

### 2.7 Discriminating individuals and speeds

To determine whether gait signatures remain characteristic to an individual across a wide range of walking speeds, we used a linear support vector machine (SVM) discrimination classification task to classify individuals based on their gait signatures across their nine speed trial conditions. To maintain consistency in the number of trials per participant analyzed, individuals with fewer than nine speed trials were excluded from this analysis. The resultant dataset comprised 14 individuals, each with nine different speed condition trials. For each data type, eight distinct SVM classifiers were trained on a progressive selection of one to eight random trials per individual over 140 total runs using built in MATLAB function ‘templateSVM’ with standardized features. The random number generator seed was set to the run number on each iteration for consistency across the different models. The average classification accuracy across the runs corresponding to the number of speed trials per individual in the training set was calculated for each data type. We conducted this analysis separately for the gait signatures across data types (Fig. 4, Discrimination task).

**Figure 4:**
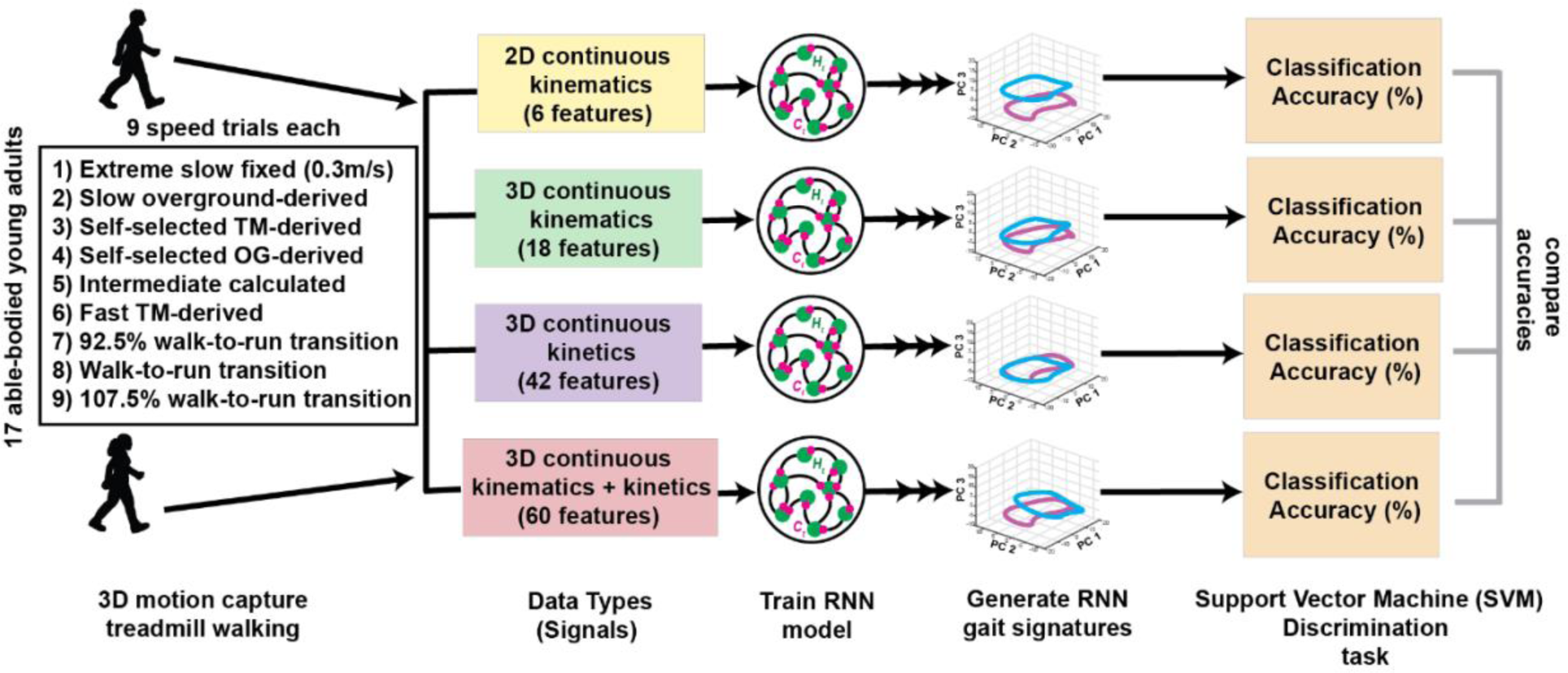
Schematic outlining the comparison of individual discriminatory power between the gait signatures generated using four different signals (2D kinematic, 3D kinematic, 3D kinetic and a combination of all signals). 3D motion capture of treadmill walking was obtained from 17 able-bodied young adults, encompassing 9 different speed conditions. The four data types were created, each with varying number of features. RNN models were trained for each data type and respective gait signatures were generated. The classification accuracy of individuals across different speeds was assessed using support vector machine (SVM) classifiers, and the classification accuracies between the four gait signature types were compared.

Because our gait signature classification accuracies do not obey gaussian statistics, we used non-parametric Mann-Whitney U tests to test for differences in classification accuracy between gait signatures generated from 2D versus 3D inputs. For classifiers trained using one to eight speed trials per participant, we compared classification accuracy and reported p-values and Mann-Whitney U effect sizes (*r*). An effect size (*r)* smaller than 0.3 indicates a small effect, a value between 0.3 and 0.5 suggests a medium effect, while an effect size greater than 0.5 indicates a large effect.

To assess the individual discriminatory potential of discrete biomechanical variables across speeds, we conducted a comparable SVM classification approach. Specifically, we focused on 26 widely recognized bilateral kinematic and kinetic discrete variables commonly used in gait analyses^[6]^, as detailed in Supplementary Table T1. Additionally, we explored the discriminatory capacity of only the 18 kinematic-only and eight kinetic-only variables in distinguishing individuals.

### 2.8 3-D Multi-dimensional scaling map to compare gait signatures

To analyze and visualize the pairwise distances between gait signatures, we employed multi-dimensional scaling (MDS) to project all gait signatures onto a single 3D gait map. This technique was used to lower the dimension of our complex high-dimensional signatures, maximally preserving relationships between individuals and trials.

To quantify the intra-vs. inter-individual differences in 3D MDS signatures, the Euclidean distances were calculated between all signatures within an individual and between individuals, respectively. Subsequently, a histogram of the Z-scored Euclidean distances was plotted to visualize both intra and inter-individual distances and the Mann-Whitney U test was performed on the distributions of distances.

### 2.9 Identifying relationships between gait signatures and speed

To determine if the relationship between 3D MDS coordinates and gait speed is linear, simple linear regression was performed separately for each of the three MDS coordinates and the speed for each individual. Box plots illustrating the slope and R^2^ values of the linear fits were generated, and the mean and 95% confidence intervals of these distributions were recorded. Additionally, p-values were recorded for the linear fits of each participant.

To test whether individuals exhibit similar linear changes in dynamics across speeds, we used linear mixed effects (LME) models to predict each trial’s 3D MDS coordinates from speed trials. We estimated positions in 3D MDS, separately for each dimension, using linear mixed-effects models with fixed intercepts and effects of speed, and a random intercept for subject. The fixed and random effects coefficients differ in each dimension. The models assume that the relationship between MDS location and trial speed is linear, while allowing for individual differences in the mean location of gait signatures in MDS space. The models aim to capture how the overall changes in gait signatures correspond to changes in speed for different subjects. MDS 3D coordinates (X, Y and Z) are the dependent variables (to be predicted), trial speed is the predictor variables and individuals’ Subject ID was used as a categorical random intercept (Equation 1).

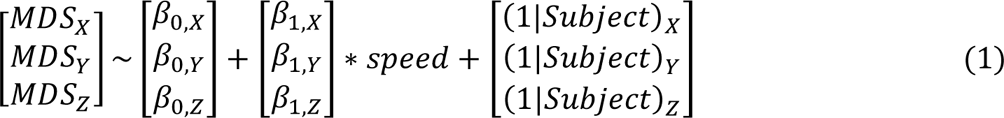

We performed a hierarchical bootstrapping analysis^[50]^ to examine how sensitive the model parameters are to variations in the input data, the number of trials chosen per individual, and the speed trials used in training. Specifically, we conducted 17 leave-one-subject-out LME models to predict 3D MDS coordinate positions from gait speed. We manipulated the number of speed trials chosen per subject (ranging from four to eight) across five iterations, with five random selections of speed trials per trial count.

### 2.10 Relationships between gait signatures and individual differences in mobility to walking speed and changes in gait signatures

To determine whether changes in gait signatures with speed are associated with individual’s functional abilities, we used a narrowing beam balance test, which has been used to characterize walking balance and motor coordination^[51]^. The narrowing beam balance score, representing the distance travelled in feet travelled along the narrowing beam, was calculated as the average distance across four trials. We used linear regression and assessed the correlation coefficient (Pearson’s r) between balance score and the following variables: a) self-selected walking speed, b) Euclidean distance between SS and extreme slow signatures and c) Euclidean distance between SS and extreme fast signatures. Additionally, to assess whether SS walking speed are associated with the extent to which participants altered their gait signatures, we performed linear regression between the Euclidean distance between SS and extreme slow signatures vs. SS speed.

### 2.11 Exploratory analysis of gait signatures and spatiotemporal variables

To determine whether changes in gait signatures with speed reflected similar findings in the literature about changes in spatiotemporal variables with speed, we conducted an exploratory analysis to determine whether changes in gait signatures (Euclidean distance) between both self-selected (SS) and extreme slow, and extreme fast speeds were associated with discrete, trial-averaged spatiotemporal biomechanical variables. Ten different Pearson’s r correlation tests were conducted for bilateral biomechanical variables: cadence, stance duration, swing duration, step width, and step length. The alpha value of 0.05 was corrected using Bonferroni method^[52]^ for the 10 different tests per speed modulation type (SS to extreme slow and SS to extreme fast) and updated to 0.005 each.

## 3. Results

In summary, individual differences in gait signatures were maintained across the full range of walking speeds from extremely slow to the walk to run transition. 3D kinematic gait signatures achieved almost perfect individual classification accuracy of 99% (CI: 99.1-100%), using four or more random speed trials in the SVM classifier training set. While individual classification performance was also relatively high for 2D kinematics (95.0%, CI: 88.1-100%) and 3D kinetics (97.8%, CI: 99.7-98.8%), it remained lower compared to that of 3D kinematic gait signatures. Linear mixed effects models revealed that individuals exhibited similar changes in their gait signatures across different speeds. This analysis uncovered a unifying framework for the impact of speed on gait signatures. We found a significant positive correlation between changes between SS and the extreme slow speed gait signatures and an external measure of balance ability (p = 0.01, r = 0.60). Furthermore, an exploratory analysis revealed that changes between SS and extreme slow signatures show significant, positive, linear relationships with speed-induced changes in two out of five spatiotemporal variables: cadence and step length.

### 3.1 Kinematics and kinetics during treadmill walking differ qualitatively across individuals and speeds

Representative right sagittal plane ankle kinematics (Fig. 1a-i) and right vertical ground reaction forces (Fig. 1b-i) were relatively consistent across all strides within an individual’s trial. Slower-speed trials resulted in fewer gait cycles than faster-speed trials over the shown 10-second period with noticeable larger joint excursions and ground reaction forces with increasing speed (Fig. 1ab-i). Despite similar general shapes of phase-averaged kinematics and kinetics across participants and speeds, there was evident nuanced variability (Fig. 1ab-ii).

### 3.2 Kinematic gait signatures demonstrate low dimensionality, contrary to kinetic gait signatures

All gait signature data types met the criterion of explaining at least 80% variance in the original data signals while using fewer than 34 PCs. However, the number of PCs required to achieve this threshold varied among different gait signature types (Fig. 3b). Specifically, six PCs of 2D kinematic signatures capture over 90% of the variance in the original sagittal plane kinematics (Fig. 3b-i). On the other hand, 3D kinematic gait signatures require 10 principal components to account for ∼83% of the variance in the original features (Fig. 3b-ii). Notably, in 3D kinetic signatures and combined kinematic and kinetic signatures, achieving at least 80% variance necessitated 29 and 33 PCs respectively (Fig. 3b-iii and iv).

### 3.3 Kinematic data types preserve individual differences in gait signatures across speeds

Phase-averaged 2D and 3D kinematic gait signatures maintain individual-specific trajectories across their entire range of walking speeds. The looped projections of the first three dominant PCs (PCs 1-3) (Fig. 5ab-i), and the second 3 dominant PCs (PCs 4-6) (Fig. 5ab-ii) belonging to each individual (specific color) cluster tightly together (Fig. 5ab-i). Higher order PCs 4-6 reveal more individual specific clustering (Fig. 5ab-ii), especially in the 3D kinematic RNN signatures (Fig. 5b-ii).

**Figure 5:**
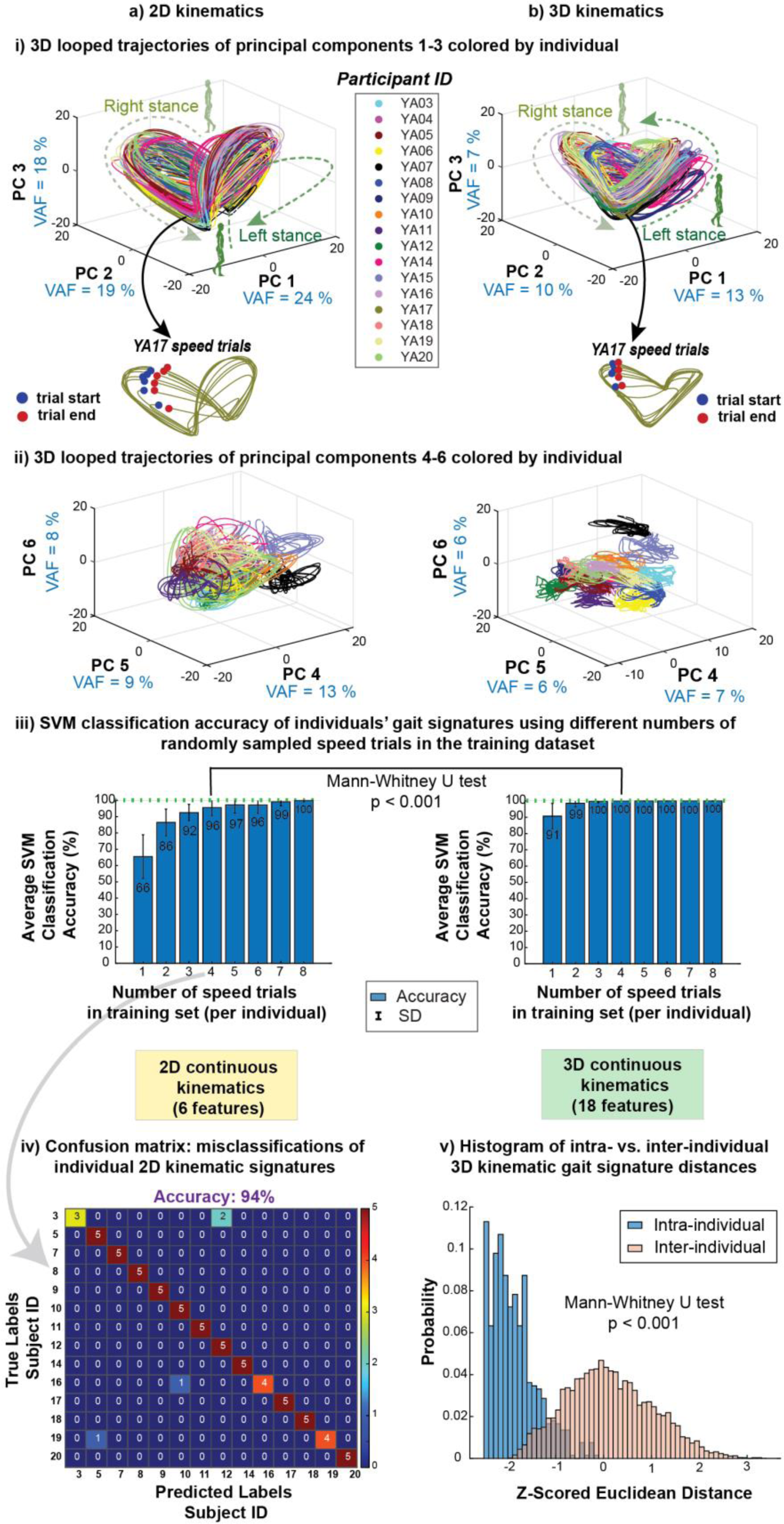
Kinematic gait signatures are individual-specific. i) 3D looped trajectories of the first 3 principal components (PCs 1-3) of a) 2D kinematic and b) 3D kinematic gait signatures are individual-specific across speeds. Individual’s trials (same color) are grouped closely together with similar shapes. ii) 3D looped representations of the second set of principal components (PCs 4-6) reveal greater individual specific clustering in a) 2D kinematic signatures, and stronger differentiation observed in b) 3D kinematic signatures. iii) Individual classification accuracy is lower using a) 2D kinematics vs. b) 3D kinematics across varied number of speed trials in the classification model training set. 3D kinematic signatures exhibited robust classification of individuals, achieving a mean accuracy of 99.8% (CI: 99.1% to 100%), with a minimum of four speed trials in the training set, surpassing the accuracy of 2D kinematics, which achieved 96% (CI: 88.2-100%). Please note that significant differences exist between 2D and 3D kinematic signature accuracies regardless of the number of speed trials in the training set. However, for clarity, we specifically emphasize the statistical comparison in the illustration using only four speeds in the training set as this is where 3D kinematic signatures attain near perfect (100%) classification accuracy. iv) A representative confusion matrix from a single classification model run shows that several individuals were misclassified when four speed trials per individual were included in the training set. v) The intra-individual trial distances in MDS space for 3D kinematic signatures are smaller than the intra-individual distances, further showcasing the individual-specificity across all speed trials of an individual.

Note that while 2D and 3D kinematic signatures were generally similarly shaped across all individuals’ speed trials, 3D kinetic signatures showed 2 subsets of signatures: some participants’ signatures tend more upwards in PC3 compared to other signatures (Supplementary Fig. S1a-i). Specifically, four participants (YA16, YA17, YA18 and YA19) appeared to have similarly shaped gait signatures compared to the other participants (Supplementary Fig. S1a-ii). This separation of signatures into 2 groups was not quite as evident in the combined 3D kinematic and 3D kinetic signatures (Supplementary Fig. S1b-i and ii). This division of 2 groupings of 3D kinetic signatures is more evident in the higher order PCs 4-6 (Supplementary Fig. S1a-iii) and 3D MDS plots (Supplementary Fig. S1ab-iv).

3D kinematic signatures consistently classified individuals with significantly higher accuracy than 2D kinematic gait signatures, regardless of the number of speed trials per individuals used in the SVM training set. The classification accuracies of 2D kinematic gait signatures were significantly lower than the 3D kinematic gait signatures across one to eight speed trials in the training set (p ≤ 0.001, -0.27 < r > -1.01). If including at least four speed trials in the training set, 3D kinematic signatures achieved almost perfect classification accuracy of 99.8% (CI: 99.1-100%) (Fig. 5b-iii), whereas 2D kinematic signatures achieved 95% (CI: 88.2-100%) (Fig. 5a-iii). To attain perfect individual classification accuracy for 3D kinematic signatures (100%), a minimum of seven training speeds are required, however, even with the inclusion of additional speed trials up to eight in the training set, 2D kinematic signatures failed to reach 100% accuracy (Fig. 5a-iii). In a representative confusion matrix of a single classification iteration using run of 2D kinematic signatures, we observed at least 3 instances of individual misclassification (Fig. 5a, iv). Classification analyses did not substantially improve when adding kinetic data (Supplementary Fig. S1a-v). Our analysis using 26 bilateral, discrete biomechanical variables (Supplementary Table T1) achieved 92.2 ± 0.1% classification accuracy with only four randomly selected speed trials per individual in the SVM training set. Kinematic-only variables yielded similar high average accuracy at 91.2 ± 0.1%. However, kinetic-only variables had significantly lower accuracy (p < 0.001) at of 64.3 ± 0.2%, with misclassifications up to 12 out of 14 individuals’ speed trials.

Analysis of the inter-and intra-individual Euclidean distances between 3D kinematic signatures in 3D MDS space demonstrates that intra-individual distances are smaller than inter-individual differences (p < 0.001), further supporting for the individual-specific nature of gait signatures within our cohort of able-bodied gait (Fig. 5b-v). The proportion of distances above two standard deviations (SD) (|z-score| > 2) were 0% and 2.2% for intra and inter-individual distances respectively. The proportion of distances below two SDs were 48.8% and 0% for intra and inter-individual distances respectively.

### 3.4 Gait signatures are modulated consistently with changes in walking speed

Despite showing that gait signatures are individual-specific across speeds (Fig. 5), 2D and 3D looped kinematic gait signatures (colored by speed) appear similar in structure and shape (Fig. 6ab-i). Both gait signature types show that slower trials (blue) are more flattened in PC3 and concentrated to the center of all speed signatures, whereas faster speeds are more expanded in PC3 and found on the outskirts of all signatures (Fig. 6ab-i, top row). This result can be seen in a representative gait signature across speeds (Fig. 6ab-i, YA11 speed trials).

3D gait maps of both 2D and 3D kinematic gait signatures colored by gait speed reveal that slower speeds (blue) across subjects are distinctly located in one region of the map and faster speeds (red) are in another region of the map (Fig. 6ab-ii). This modification by speed is more noticeable in 2D signatures than in the 3D signatures.

We found that across participants, the 3D MDS representation of gait signatures change in a similar direction with changes in gait speed (Fig. 6ab-iii). Notably, even at extremely slow speeds (0.3 m/s), similar to those observed in stroke survivors, the gait signatures maintained a consistent linear relationship with faster speeds. Linear mixed effects models accurately explained the relationship between individuals’ MDS coordinate representations of their 3D kinematic gait signatures and speed with high accuracy (R^2^ > 0.95) for each coordinate (Fig. 6b-iii). However, the accuracy was lower for 2D kinematics (R^2^ > 0.89) (Fig. 6a-iii). Note that the p-value for the linear MDS coordinate fits in the Y axis of the 3D kinematic signatures were not significant (p = 0.28).

**Figure 6:**
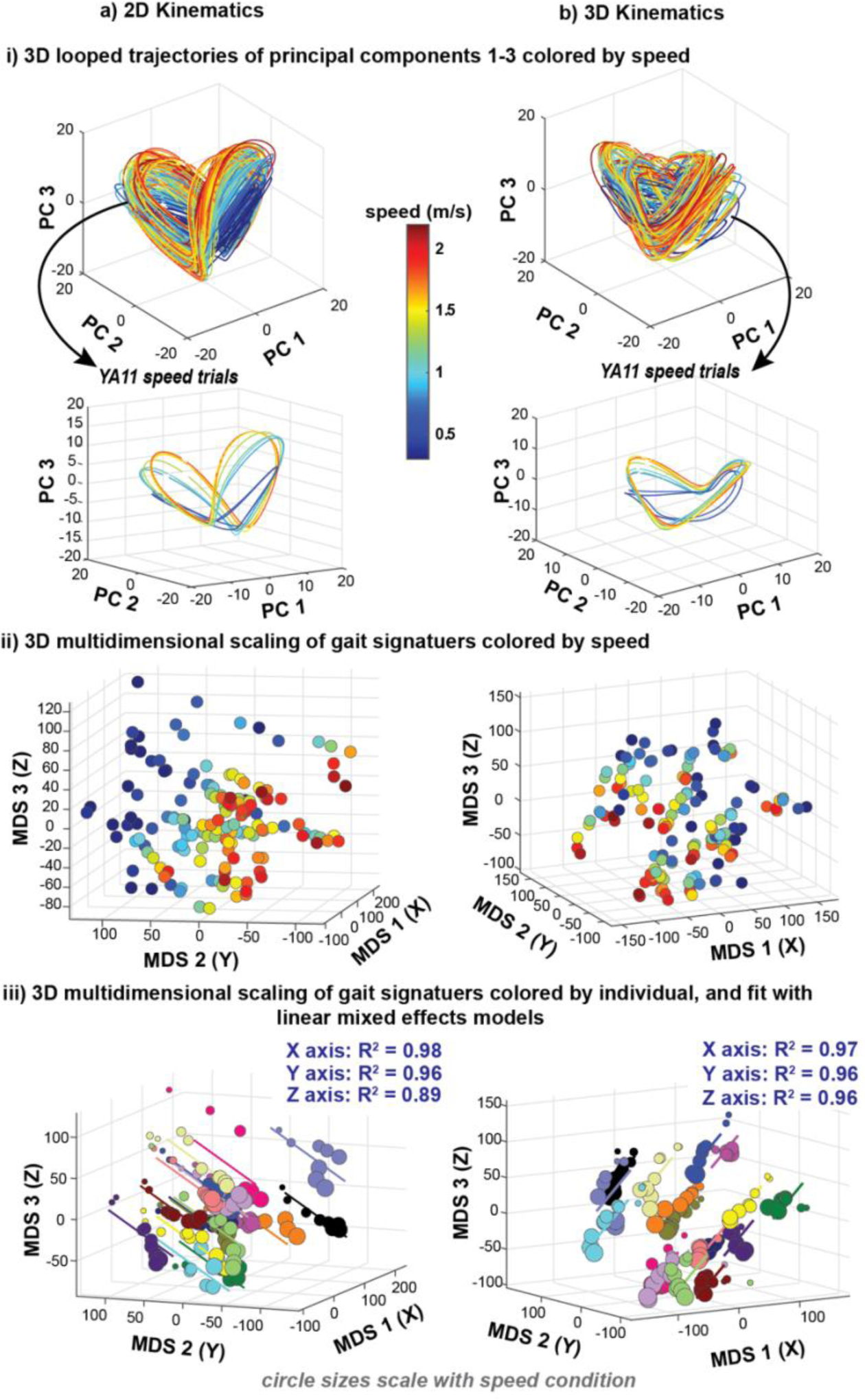
Gait signatures hold information about walking speed. i) 3D looped trajectories of principal components 1-3 colored by gait speed show that similar speed trials are shaped similarly in a) 2D kinematic and b) 3D kinematic signatures. Slower speed signatures (blue) are concentrated in the center of all signatures and faster speed signatures (red) on the outskirts, fanning outward and upward in PC3. ii) 3D MDS visualizations of all signatures colored by speed illustrates that slower speeds (blue) across individuals are in the top left region of the map for a) 2D kinematic signatures and in the right half region of the gait map for b) 3D kinematic signatures. iii) 3D MDS visualizations of all signatures colored by individual fit with linear mixed effects models show that individuals gait signatures change similarly and linearly with change in gait speed.

### 3.5 Within individuals, 3D MDS coordinates of gait signatures vary linearly with speed

Linear regression of each individual’s 3D MDS gait signature coordinates separately (XYZ) showed consistently strong and significant associations between changes in the X-axis (R^2^ > 0.50; p < 0.02) across participants, but not in the Y-or Z-axes (Fig. 7a). X and Z significant slopes are negative, whereas Y has significant slopes that are positive and negative (Fig. 7b). Moreover, distinct clusters of individuals exhibited similar changes in their gait signatures with speed were evident, particularly in the X and Z coordinate slope values. These clusters were observed: a) around the mean value (red dotted line), b) above the upper limit of the 95% confidence interval (gray dotted line), and c) below the lower limit of the 95% confidence interval (Fig. 7a).

**Figure 7.**
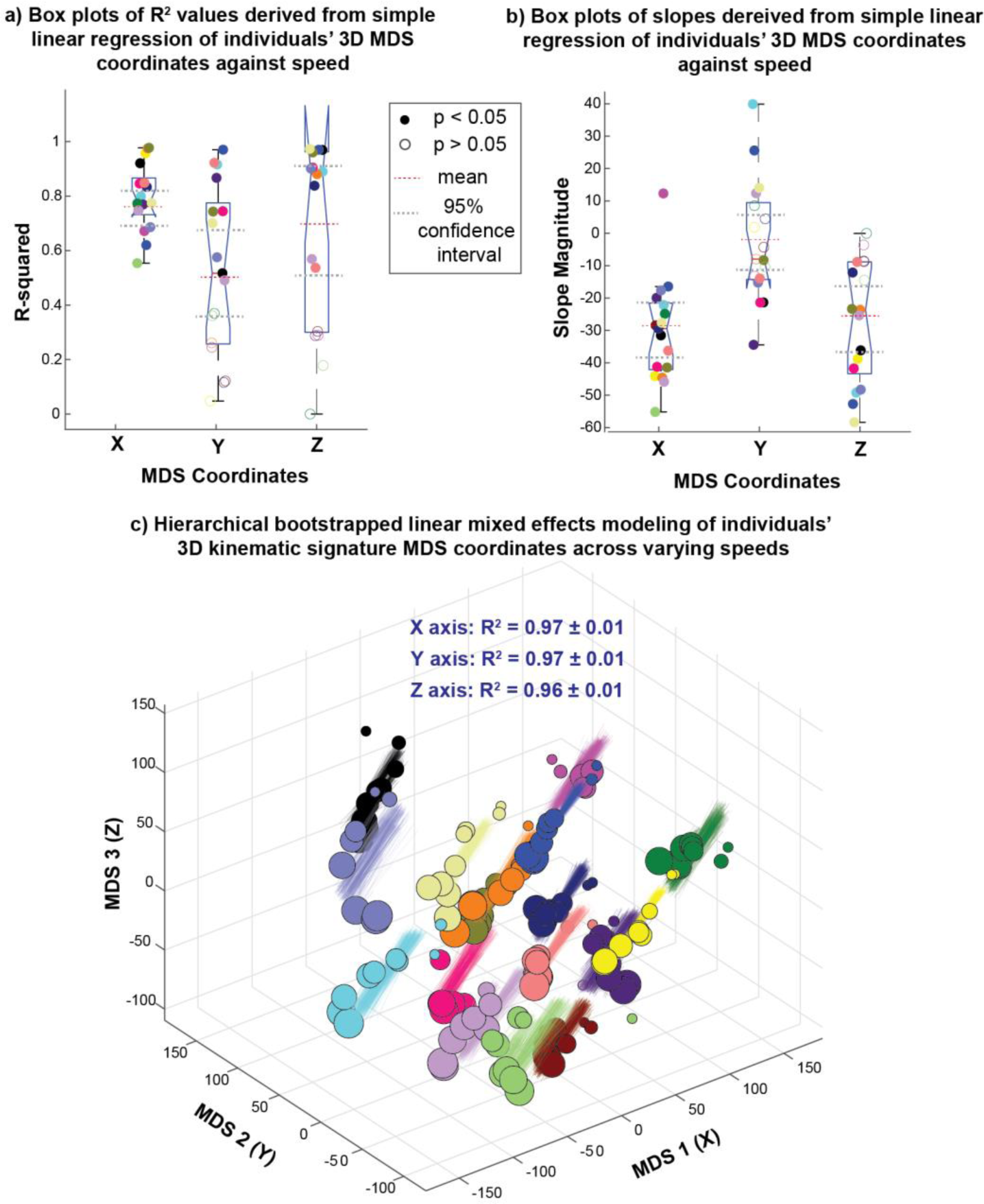
Individual’s gait signatures change linearly with speed. Simple linear regression of individuals’ 3D MDS coordinates vs. speed show similar a) R^2^ values and b) slopes across individuals. c) Hierarchical bootstrapping of linear mixed effects shows that the linear relationship of MDS coordinates with speed is robust across variability in model input data, the number of speed trials selected per individual, and the randomness of the selected speed trials used in model.

### 3.6 Stability of linear mixed effects model parameters across individuals

Linear mixed effects (LME) models tested the similarity of the linear relationships between gait signatures and speed across the entire cohort. Hierarchical bootstrapping results confirmed that robustness of the LME models to variations in input data, the number of trials per individual and speed trials used to train the model. Consequently, LME models were deemed suitable for accurately predicting the locations of 3D kinematic gait signatures with respect to speed (Fig. 7c). Note that a few Y-coordinate models resulted in non-significant p-values. The X, Y, Z sensitivity of the B_1_ (fixed slope) parameter was -0.30 ± 0.03, -0.02 ± 0.02 and -0.27 ± 0.03 respectively. The X, Y, Z sensitivity of the B_0_ (fixed intercept) parameter was 37.5 ± 5.7, 3.7 ± 5.8 and 35.0 ± 5.5 respectively. The standard deviation ranges of individual random intercepts for the X, Y, Z coordinates were [4.3 – 9.7], [4.2 – 8.2] and [3.9 – 6.4] respectively.

Histograms of residuals across the three LME models (representing X,Y,Z coordinate locations) exhibited normal distributions centered around zero (Supplementary Fig. S2a). Furthermore, residual vs. predicted coordinate plots show that the variance of residuals across various predictions are constant (homoscedasticity) (Supplementary Fig. S2b), meaning that the models are generally well-behaved.

### 3.7 Associations between gait signature changes and balance ability

Walking balance ability may be associated with the extent to which individuals alter their gait signatures across speeds. We identified a moderately positive correlation between individuals’ narrowing balance beam score and changes in gait signatures (Euclidean distance between self-selected and extremely slow speed walking gait signatures) (p = 0.01, r = 0.60) (Fig. 8a). Individuals with better balance (higher scores) alter their gait signatures more from SS to extreme slow. Additionally, we observed a moderate positive correlation between participants’ SS speed and their performance on the narrowing balance beam (Fig. 8b), where participants’ with faster SS speeds exhibited better performance compared to those with slower SS speeds (p = 0.02, r = 0.57). Further, individuals with slower SS speeds changed their gait signatures less between the SS and extreme slow speeds (Fig. 8c). However, changes in gait signatures between participants’ SS and extreme fast speeds were not associated balance (p = 0.15, r = -0.37) (Supplementary Fig. S3a), or self-selected walking speed (p = 0.07, r = -0.45) (Supplementary Fig. S3b). In a 3D MDS map of all gait signatures, there is no discernable trend in the spatial arrangement of individuals’ according to their narrowing balance beam score (where similar colors indicate similar balance scores) (Fig. 8d). For instance, the 2 groups of gait signatures colored orange are located distant from each other, and both are close to individuals with significantly lower narrowing balance beam scores (Fig. 8d).

**Figure 8:**
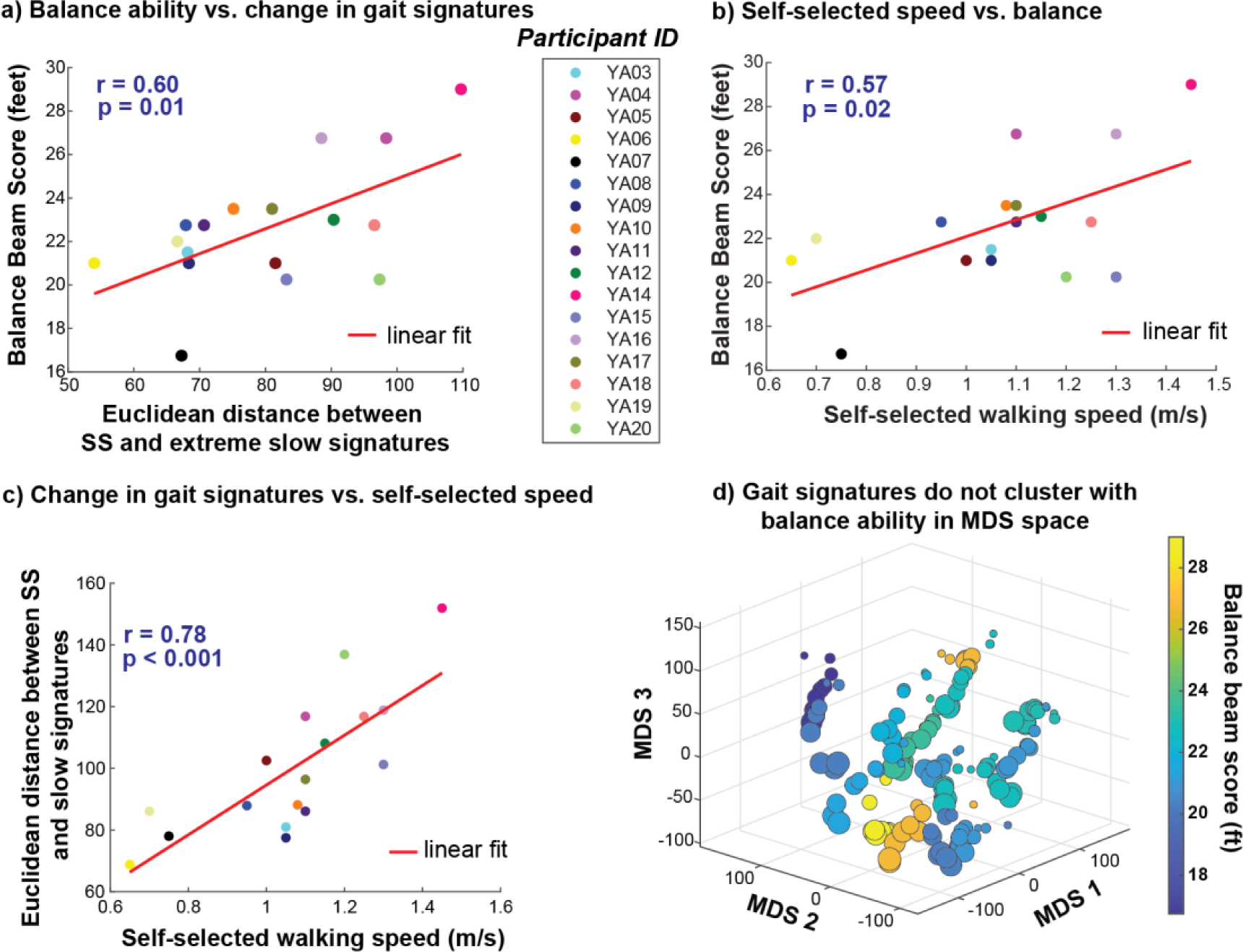
Balance ability may be associated with the extent to which individuals modulate their gait signatures with changes in speed. a) A moderately positive linear relationship exists between balance ability and the change (Euclidean distance) between SS and extreme slow speed gait signatures-individuals with better balance modulate their gait signatures more when reducing speed from SS to extreme slow walking speeds. b) A moderately positive linear relationship exists between self-selected walking speed and balance ability-individuals with better balance prefer walking at faster self-selected walking speeds. c) A strong positive linear relationship exists between change in gait signatures between SS and extreme slow speed and self-selected walking speed. d) 3D MDS representation of all individuals’ gait signatures colored by their narrowing balance beam score reveal no clustering of signatures in this space.

### 3.8 Exploring associations between changes in gait signatures and discrete spatiotemporal variables with speed

Changes in gait signatures from SS to extreme slow speeds, but not extreme fast speeds, are correlated with changes in two of five spatiotemporal biomechanical metrics (cadence, step length, swing duration, stance duration and step width). Significant correlations were identified between SS and extreme slow speed gait signatures and changes in right cadence (p = 0.002, r = 0.68) (Fig. 9a) and left step length (p = 0.001, r = 0.72) (Fig. 9b). However, there was no significant correlation between any of the 5 bilateral spatiotemporal variables and changes in gait signatures between SS and extremely fast walking speeds (p > α_Bonferroni_ = 0.005).

**Figure 9:**
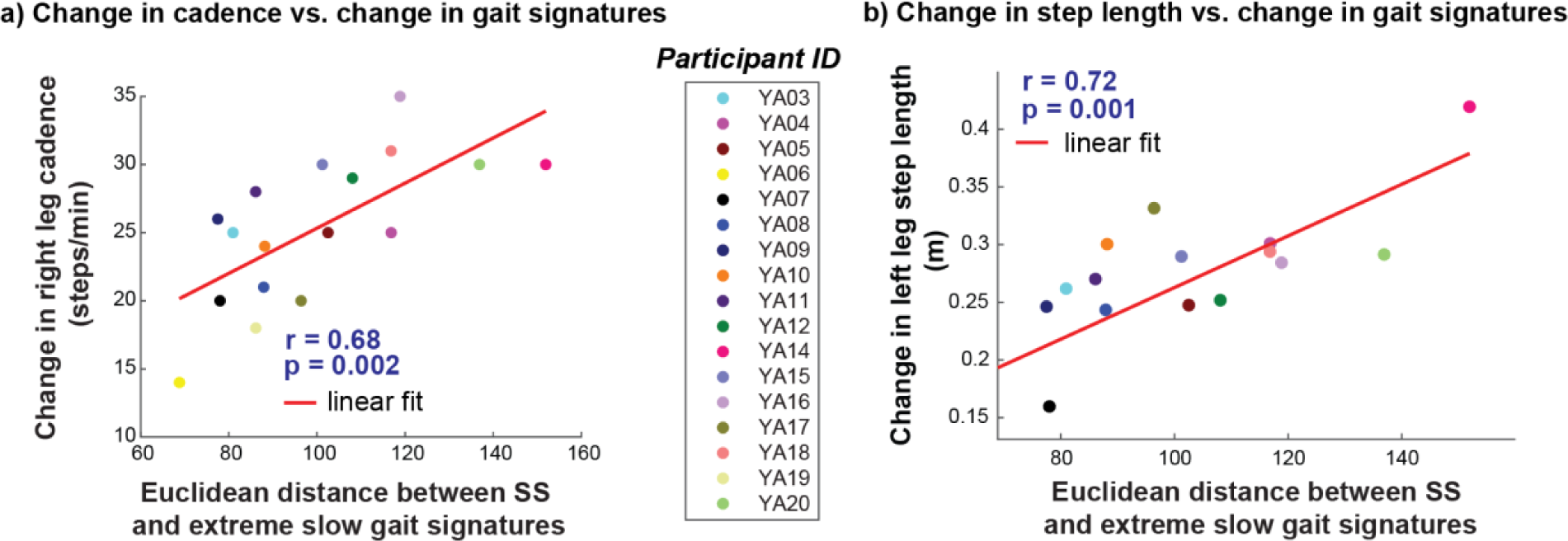
A strong positive linear relationship exists between the Euclidean distance between self-selected and extreme slow speed signatures and changes in a) right leg cadence and b) left leg step length

### 3.9 Discrete spatiotemporal variables show strong linear relationships speed, but cannot distinguish between individuals

Trial-averaged discrete variables also showed linear relationships with gait speed (Supplementary Fig. S4). Cadence and step length have strong, positive relationships with increasing gait speed (Supplementary Fig. S4a-i and iii), while swing duration and stance duration show strong, negative relationships with increasing speed (Supplementary Fig. S4a-ii and iv) at the Bonferroni-adjusted significance level of α = 0.005 for n = 10 variables (all p-values < 0.001) (Supplementary Fig. S4a). The variability of these variables at the extreme slow speed (0.3m/s) showed high variability (Supplementary Fig. S4). Stance width did not demonstrate a significant correlation with speed (p = 0.9, r = 0.03). Spatiotemporal variables only discriminated individuals with average classification accuracy of 55.7 ± 0.2%. A confusion matrix of a representative classification run illustrates that as many as 11 out of 14 individuals were misclassified (Supplementary Fig. S4b).

## 4 Discussion

### Summary

Machine-learning-based dynamic gait signatures holistically encode individual differences and systematic changes in gait dynamics with walking speed. Individual differences in gait amongst able-bodied young adults are distinguishable within a common, low-dimensional latent space of the gait dynamics model regardless of differences in how slow or fast they walk. Differences across individuals are evident using RNN models based on both 2D and 3D kinematics, suggesting that consistent gait coordination patterns within individuals can be captured from camera-based measures. In contrast, gait signatures trained using kinetics did not improve discrimination and their representations were high-dimensional, suggesting that kinetic variables do not maintain consistent coordination across speeds. Although there are unique differences between individuals’ gait signatures, the changes in these signatures as speeds change are predictable, enabling us to identify a unifying linear relationship that explains how able-bodied individuals’ gait signatures alters with speed. Within this common linear relationship, however, the degree to which individuals modify their gait signatures is variable and is related to their balance ability and self-selected walking speed. Overall, these insights underscore the need to identify unifying principles regarding the physiological or biomechanical factors that underpin these changes. The gait signatures approach may be useful to identify individual differences in gait across a variety of applications, notably in sports and personalized gait rehabilitation.

The individual-specific gait signatures discovered by our framework are consistent with the idea that individuals maintain consistent, highly characteristic gait dynamics across a variety of walking speeds^[3,4,53]^. Our study complements previous research by showing that individuals’ gaits remain individual-specific across a wide range of speeds, rather than solely at self-selected walking speeds^[1]^ or a more limited range of speeds^[6]^. Our cohort of able-bodied adults served as a rigorous test case, demonstrating the robustness of individual classification even in a healthy, young population with relatively similar dynamics. These results suggest potential discriminatory effectiveness in stroke and other patient populations as well. We show that even at very slow speeds, able-bodied gait signatures can be approximated by a linear model. Notably, prior studies have shown that walking at extremely slow speeds alter baseline gait coordination more profoundly than fast walking speeds^[40,41]^, and in our dataset, we observe high inter-individual variability in spatiotemporal parameters at very slow walking speeds. We thus postulate that walking at extreme slow speeds may necessitate distinct gait dynamics, given that observable differences in muscle coordination also exist at slow versus self-selected walking speed^[54]^. This effect could be attributed to the fact that slow walking is more akin to a postural task involving a series of weight shifts. Further, we hypothesize that there may be a greater deviation in dynamics near individuals’ walk-to-run transition speed. However, our gait signatures remained individual-specific and approximately linear across the entire spectrum of walking speeds, even at extreme fast and slow speeds. Capturing the processes governing the progression of inter and intra-limb gait variables over time offers a more comprehensive speed-independent characterization of individuals.

Kinematic data adequately capture individual differences in gait dynamics that differentiate individuals. Here we compared 3D kinematic signatures to the 2D kinematics approach from our prior work^[6]^. We found a small but significant improvement in individual discrimination using 3D kinematic signatures. However, the 96% and higher average discriminatory accuracy of 2D kinematics suggests a significant amount of information can be garnered solely from motions in the sagittal plane. Improvements in the individual classification accuracy by 3D kinematic gait signatures are likely due to the added information from movement in the frontal and transverse planes. For patient populations with more frontal-and transverse-plane gait deviations^[23–25]^, generating gait signatures from 3D kinematic data may be more important for differentiating between individuals. Additionally, the choice between 2D and 3D kinematics affects equipment requirements, with 3D analysis often requiring multiple cameras compared to 2D analyses^[55–57]^. The efficacy of using 2D kinematics in gait analysis has already shown promise in conditions such as spastic tertraparesis^[58]^, cerebral palsy^[59]^, and cancer survivors^[60]^. Moreover, 2D kinematics-based gait signatures, such as from gait videos, may still enable characterization of gait dynamics in diverse cohorts when 3D motion capture is not feasible^[61–63]^.

We were surprised that the addition of kinetics did not improve the individual distinguishability of gait signatures. The fact that kinetic data could not be modeled in a low-dimensional space within the model suggest that the dynamics governing kinematics are sufficient to distinguish individuals across speeds compared to the dynamics of forces, which are more directly related to the biomechanical demands of maintaining dynamics underlying kinematic patterns. Considering that the recurrent neural network (RNN) learns to model the evolving neuromechanical dynamics of gait over time, it is plausible that the RNN may be able to encode similar information about an individual to what is contained in kinetic data. Moreover, previous research has used kinematic data to infer kinetic data^[29,30]^, suggesting that kinematics encompass information integral to kinetics. Whether kinetic data enhance classification accuracy in people with neuro-pathologies such as stroke needs more evaluation in future work. Nonetheless, eliminating the need for costly force plates in gait analysis remains advantageous.

Our data show that able-bodied gait signatures change predictably with speed. Despite individual-specific differences in gait signatures, we found consistent directional changes in 3D MDS representations of gait signatures with speed across participants. Similarly, many researchers have found linear relationships between simple discrete spatiotemporal variables with gait speed^[14,16,17,36,37,64]^. These variables, however, describe aggregated information across individuals’ entire stride (e.g., peak, or averaged gait metrics). We show that changes in our gait signatures with speed are indeed moderately correlated with spatiotemporal variables such as cadence, step length, stance duration and swing duration, suggesting that our gait signatures framework holistically captures the information found in these composite variables. Furthermore, studies have shown significant relationships between gait speed and discrete kinematic^[39,65–67]^ and kinetic variables^[39,68]^. However, these metrics only provide information at discrete points in the gait cycle, disregarding potentially meaningful information at other time points during the gait cycle. We showed that gait signatures classified individuals with higher average classification accuracy (∼99%) than spatiotemporal variables (∼56%), discrete kinematic (∼91%), and discrete kinetic (∼64%) variables. Given that our framework uses continuous, synchronous, multi-joint data to construct gait signatures, our current results provide further support that our approach provides a more comprehensive measure of individuals’ gait coordination over time.

Aspects of individual differences in mobility– even in able-bodied young adults– appear to play a role in shaping individuals’ gait dynamics and the ability to adjust them across speeds. First, we observed that baseline gait dynamics undergo more substantial changes with extreme slow walking speeds compared to extreme fast speeds. Our results suggest that individuals with better balance may more flexibly modulate their gait dynamics from SS to extreme slow walking. Second, correlation analyses suggested that participants who preferred faster self-selected walking speeds tended to perform better on the narrowing balance beam. We infer that better balance and the extent to which individuals’ modulate their gait signatures likely emerge from a common underlying construct (a confounding variable)-walking speed. Slower walking speeds are associated with worse balance and greater risk of falls, particularly among older adults and individuals with neurological impairment^[69,70]^. Thus, the habitual walking speed preferences observed among able-bodied young adults may be linked to their individual balance capabilities.

The inclusion of extremely slow speeds in our analysis is particularly relevant for comparisons between able-bodied individuals and impaired populations who may walk at very slow speeds. In our previous study^[6]^, we investigated both able-bodied and stroke individuals across a narrower speed range, ranging from their self-selected speeds to their comfortable fast speeds, revealing minimal overlap between the speeds of the two cohorts. The speed limitation stemmed from neuromechanical impairments and safety concerns for stroke survivors. Specifically, stroke survivors walked from 0.3m/s to 0.9 m/s, while able-bodied individuals ranged from 0.9 to 1.6m/s^[6]^. Thus, the incorporation of these much slower speed signatures for able-bodied young adults in our current work facilitates a more unbiased comparison of gait dynamics for impaired populations, such as stroke survivors. Conversely, the inclusion of speeds up to the walk-run transition may facilitate comparisons with cohorts comprising individuals who walk at exceptionally fast speeds, such as competitive speed walkers.

It is important to address some potential biases and limitations in our methodology. First, we enforced a fixed extreme slow speed of 0.3m/s across participants to ensure the speed reflects that of impaired gait speeds such as stroke survivors. This imposition may have introduced a floor effect, particularly affecting taller individuals or those with longer legs. Future studies should consider scaling speeds to leg length^[71]^. Additionally, we did not use a multi-session approach or have a rigorous practice session^[72]^. Participants may select different transition speeds given more time on the treadmill. However, given the linear regression results, we do not expect conclusions to change at slightly higher or lower transition speeds. Kinematic and kinetic variables in the frontal and rotational planes may be prone to measurement errors and greater inter-individual variability due to marker positioning, but the evaluation of within-individual changes across speeds reduces this concern in the current study^[27,73–75]^. To improve visualization of our data and simplify analyses, we employed 3D multi-dimensional scaling of our high dimensional gait signatures (> 1000D), which may have resulted in a loss of information about relative similarity between individuals and trials. The results may have changes if a higher-dimensional representation of dissimilarity was used. Our use of support vector machine classifiers to distinguish individuals across speeds presents interpretation challenges, as the learned decision boundary may be complex and difficult to interpret, providing limited insight into the underlying relationships driving classification outcomes. Additionally, using a linear mixed effects model on a relatively small dataset raises concerns regarding the reliability of estimates for the variance components for random effects. These random effects capture variability among the individuals, potentially leading to a reduction in residual variability and an inflated perception of explained variability, thus inflating the R^2^ values.

The results of our study hold significant implications for real-world applications, particularly in gait research, sports training, and gait rehabilitation. Our work can potentially aid researchers with determining the minimum equipment required for constructing gait signatures capable of effectively characterizing individual differences in gait dynamics. The ability of gait signatures to reliably identify individuals walking at any gait speed may also be useful in research where biometric recognition using machine learning is important^[76,77]^. The finding that 3D MDS gait signatures can be reasonably predicted from a limited set of speeds may offer a valuable framework similar to velocity-based training in sports conditioning^[78,79]^, potentially enabling trainers to pinpoint optimal training speeds for athletes or prescribe training intensities to induce desired changes in in movement quality. The linear relationship between gait signatures and speed could inform modeling and control applications, aiding the development of more efficient locomotion strategies for human-robotic systems^[80]^. This relationship is particularly relevant given that previous systems typically operated at speeds equivalent to or lower than a participant’s comfortable overground speed^[81,82]^. For instance, adaptive prosthetic devices and exoskeletons could be designed to mimic an individual’s natural gait patterns more closely across a range of speeds. However, while gait signatures generally change linearly with speed in able-bodied adults, this relationship may not be the case for impaired individuals, underscoring the importance of future studies in understanding individual-specific response to conditions and interventions. Moreover, individual-specific signatures hold clinical value by extending treatment monitoring and measurement beyond discrete clinical measures^[83]^. Gait signatures may also facilitate the design of personalized gait rehabilitation programs by leveraging insights into how individual gait characteristics predict treatment efficacy^[84]^. The direct association between changes in gait signatures and improvements in gait quality remains unverified. Future work should explore the relationship between gait signatures and gait quality, as well as define clinically meaningful changes. Nonetheless, more research is required to understand how impaired gait signatures change with speed, considering safety concerns and the capabilities of impaired populations. Our study opens new avenues for personalized rehabilitation interventions and enhanced sports performance through informed speed selection.

## 5 Conclusions

We used our previously developed gait signatures framework to show that gait dynamics remain individual-specific across a wide range of speeds, suggesting that this framework may be useful in characterizing individual differences in gait impairments and inform training or rehabilitation personalization. Approximately linear changes in able-bodied young adult gait signatures with gait speed allows inferences to be made about their gait dynamics at unmeasured speeds. Further, changes in gait signatures from self-selected to extreme slow speeds were correlated with balance ability, individuals’ self-selected walking speed, and discrete spatiotemporal variables, pointing to specific factors that may be shaping changes in dynamics with speed. While this work focuses on solely able-bodied individuals, future work should include impaired cohorts before similar gait analyses can be translated to clinical practice. By considering the dynamic evolution of multiple gait variables over time and their modulation with speed, researchers and practitioners can gain a deeper understanding of how individuals’ gait patterns adapt across different speeds. This perspective can enable the development of interventions tailored to meet the specific needs of individuals with gait impairments.

## 6 Supplementary Figures

**Supplementary Fig. S1:**
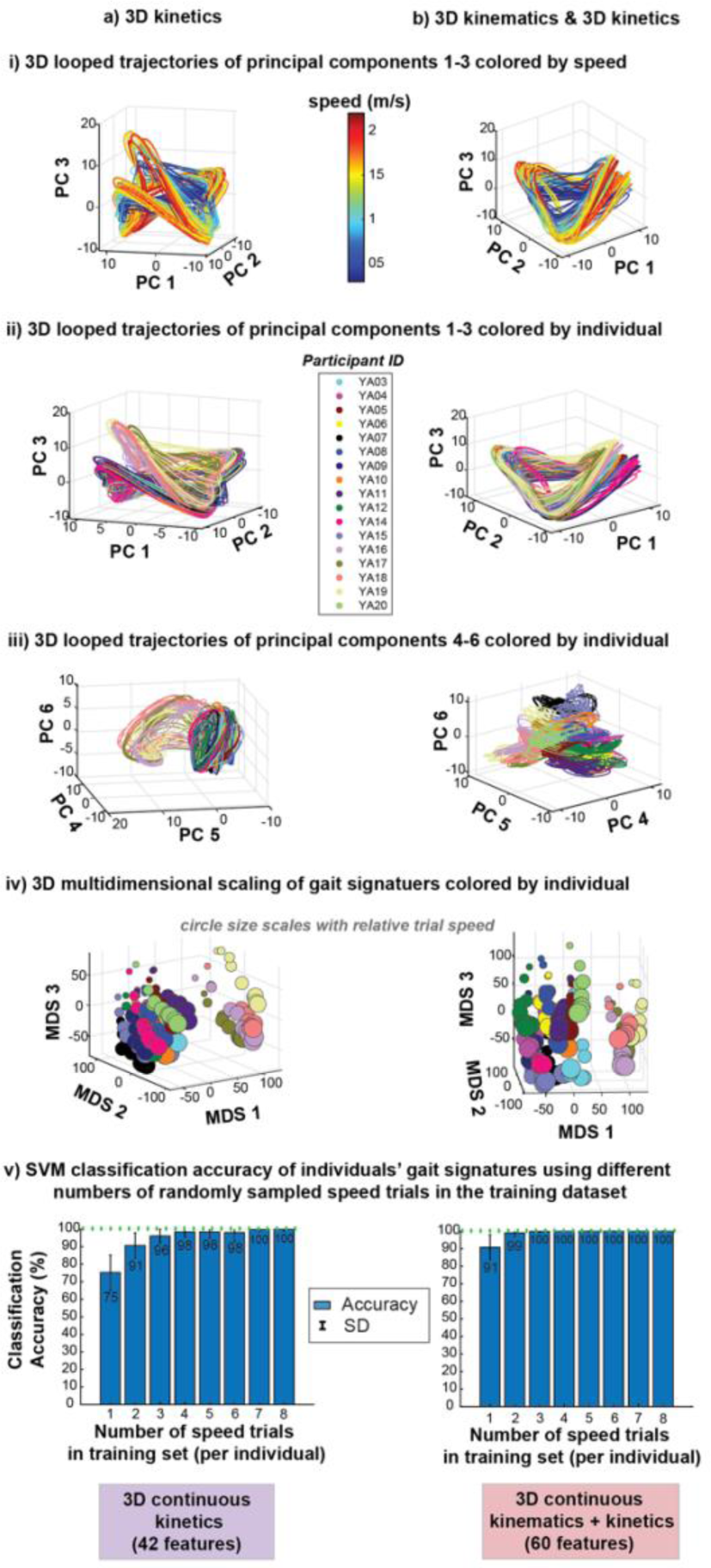
Visualization of kinetic-based gait signatures a) 3D kinetics and b) 3D kinematics & 3D kinetics. i) 3D looped trajectories of the first 3 principal components (PCs 1-3) of the gait signatures colored by speed shows that gait signatures at faster speeds (red) were more expansive than those at slower speeds (blue) ii) 3D looped representations of the first 3 PCs 1-3 of the gait signatures colored by individuals revealed that signatures are individual-specific across speeds. Kinetic signatures (a) appeared to form 2 groups of individuals with differing looped trajectories. iii) 3D looped representations of the second set of PCs 4-6 of the gait signatures colored by individual showcased individual-specific signatures. iv) 3D MDS visualizations of all signatures colored by individual further reveals a splitting of individuals into two groups. v) Individual classification accuracy was relatively high using a) 3D kinetics and b) 3D kinetics & kinematics across varied number of speed trials in the classification model training set.

**Supplementary Fig. S2:**
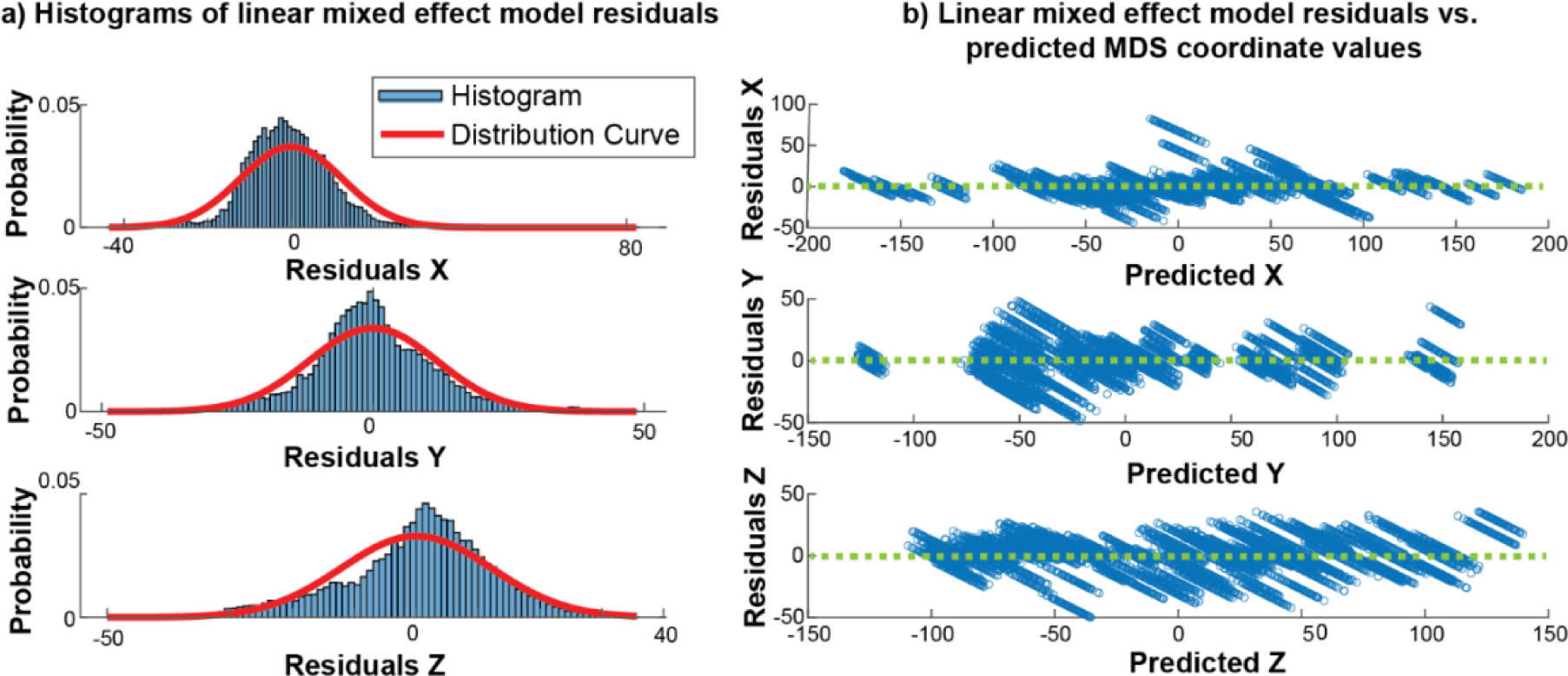
Evaluation of LME model fits. a) Histogram of residuals across 3D coordinate LME models are centered around zero. b) Residuals vs. predicted values reveal homoscedasticity (fluctuation around zero) regardless of prediction value.

**Supplementary Fig. S3:**
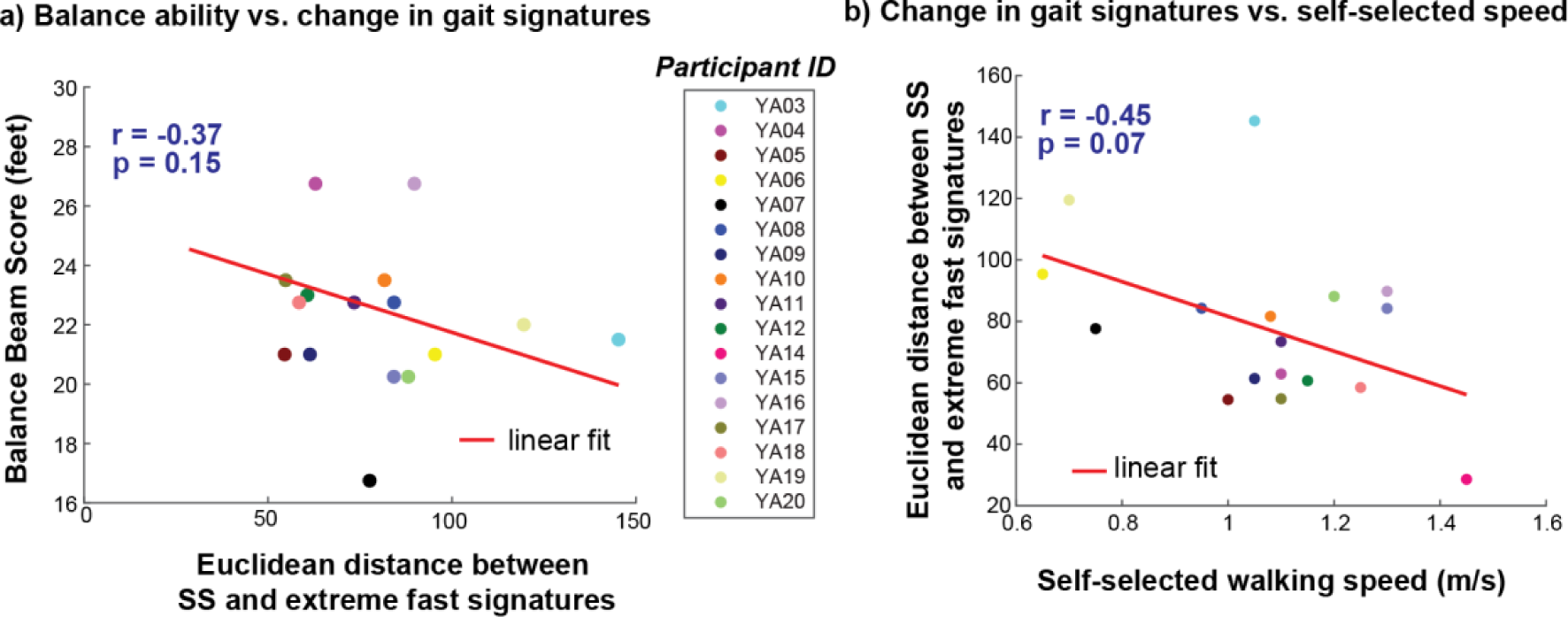
Correlation plots showing no significant linear relationships between Euclidean distance between SS and extreme fast (walk to run transition) speed signatures and a) narrowing balance beam score and b) self-selected walking speed.

**Supplementary Fig. S4:**
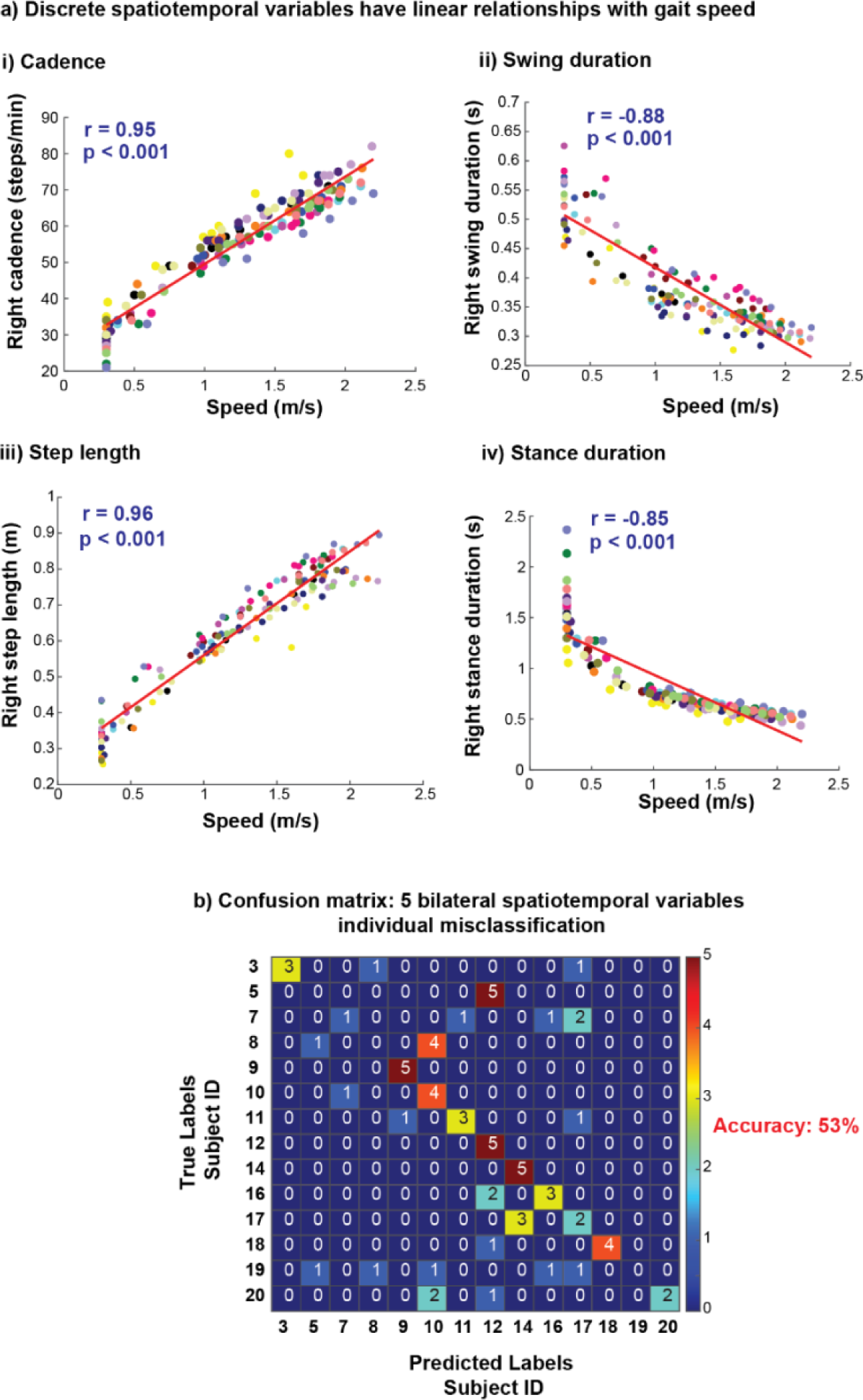
Discrete biomechanical variables show strong, linear relationships with speed. a) Discrete spatiotemporal variables i) cadence and iii) step length show strong positive linear relationships with increasing gait speed and variables ii) swing duration and iv) stance duration show strong negative linear relationships with increasing gait speed. b) Five bilateral spatiotemporal discrete variables (cadence, step length, swing duration, stance duration and step width) were unable to classify individuals with high accuracy (53%). A confusion matrix, derived from a single run of a linear support vector machine classification model, illustrates that multiple individuals were misclassified.

**Supplementary Table. T1:** 13 commonly used discrete biomechanical variables assessed bilaterally in gait analysis.

| Kinematics | Kinetics |
| --- | --- |
| Step length | Peak anterior ground reaction force |
| Peak trailing limb angle | Push off integral |
| Peak hip hike | Peak ankle moment |
| Double support duration | Peak ankle power |
| Stance duration |  |
| Swing duration |  |
| Ankle angle at heel strike |  |
| Ankle angle at toe off |  |
| Knee angle at midstance |  |

## 7 Data Availability Statement

All data and code that support the findings in this paper has been deposited at GitHub: https://github.com/bermanlabemory/GaitSignatures_HealthyYoungAdultStudy. The RNN model training and gait signature development was conducted in Python programming language. The data analysis of the generated gait signatures was conducted in MATLAB 2023a (MathWorks). The deposited materials are accessible to enhance reproducibility and advocate for open science. Additional data supporting this study’s findings are available on request from the corresponding author, Taniel Winner.

## 10 Author contributions

T.S.W, T.M.K, L.H.T, and G.J.B contributed to the conception and design of the work. T.S.W, M.C.R, and T.M.K contributed to the data collection. T.S.W. contributed to code development, analysis, generation of results and developed the initial draft of the manuscript. All authors contributed to the analysis, interpretation of the results, the writing and revision of the manuscript.

## Acknowledgements

We would like to express our gratitude to Alexandra Slusarenko for her assistance with experimental data collection. We thank Benjamin Fargnoli and Jifei Xiao for their assistance with Vicon data processing. Additionally, we thank the participants who generously volunteered their time for this study.

